# The evolution of cooperation under local regulation and non-additive gene action: building on Hamilton’s ideas

**DOI:** 10.1101/007682

**Authors:** Roberto H. Schonmann, Robert Boyd, Renato Vicente

**Affiliations:** Dept. of Mathematics, University of California at Los Angeles, CA 90095, USA; School of Human Evolution and Social Change, Arizona State University, Tempe AZ; Santa Fe Institute, Santa Fe NM; Dept. of Applied Mathematics, Instituto de Matemática e Estatística, Universidade de São Paulo, 05508-090, São Paulo-SP, Brazil

**Keywords:** Cooperation, population viscosity, Hamilton’s rule, group-structured populations, islands model, population regulation, non-linear marginal fitness function

## Abstract

We study evolution of cooperation in a population structured in a large number of groups of variable size, connected by random migration at rate *m*. Social interactions, including cooperation and competition occur only inside the groups. Assuming that groups are large, we define a parameter *λ* that measures the strength of the local regulation, i.e., the rigidity of group sizes. Individuals are of two possible genotypes, one typically assumed to produce a non-cooperative phenotype and the other a phenotype that is cooperative with all members of its own group. Gene action may be additive, producing fitness functions that are linear in the number of cooperators in a group, or not. Assuming weak selection, we obtain the following two contrasting conclusions. (1) “Hamilton regime”: If *λ << m*, then cooperative behavior can spread under a certain condition, which in the additive, i.e., linear, case is precisely Hamilton’s rule. The general version of this condition is also relatively easy to apply and is based on Wright’s classical beta distribution for the frequency of alleles in infinite island models. We call it the “beta version of Hamilton’s rule”. (2) “Taylor regime”: If *m << λ*, then cooperation that is costly to the actor is eliminated by selection.

## 1 Introduction

One of Hamilton’s pivotal contributions to our understanding of how natural selection can lead to the spread of cooperative behavior was the observation that population viscosity could lead to the type of gene assortment needed for this to happen [12]. The idea is that an individual’s location throughout its life is correlated with its place of birth. And hence, even if cooperators provide benefits to everyone in their neighborhood, the benefits would flow to individuals who are more strongly related to the cooperator than an average member of the whole population.

It was nevertheless soon realized [13, 10, 14] that this idea may not work, since individuals also compete more with their neighbors than with an average member of the population, and hence the benefits of cooperation would be counteracted by this competition. Population viscosity generates relatedness among neighbors, but the benefits from cooperation must be compared to the opposite effects of local competition. This issue was made salient in two papers: simulations reported in [37] and a theoretical work presented in [32]. The latter assumes an island population structure, and a life-cycle in which groups produce a large number of offspring; a fraction *m* of these emigrate and the rest remain in their natal group; these juveniles then compete locally for the *n* spots available in each group. Taylor [32] showed that the effects of cooperation and local competition cancel out, so that only acts that produce a net benefit to the agent itself can be selected.

It was noted however, in e.g., [13, 32, 33], and elaborated in, e.g., [9, 22, 34, 17, 8, 1, 25, 19], that incorporating elasticity in group size, or group competition can allow cooperation to spread. Those papers provided mechanisms for viscosity to promote the evolution of cooperation, but did not fully clarify what parameters and forces should be compared in deciding when it could happen. It is worth also pointing out that the inclusion of elasticity into the models is usually considered a delicate affair (see, e.g., [8], p. 1714).

The effect of non-additive gene action on the spread of cooperation is another important unresolved issue. Most of the literature assumes that when selection is weak, the benefits of group cooperation on an individual are a linear function of the number of cooperative individuals in its social environment; in other words, that gene action across individuals is additive. (See [28] for a detailed discussion.) Additivity of gene action across individuals is a very special assumption, that is inconsistent with some data [5, 31], and fails when cooperative behavior is contingent on the past behavior of others in the group, as for instance in the iterated public goods game studied in [16, 2, 29].

Motivated by these two issues, here we analyze a model in which the amount of local competition can vary and that we call “islands with local regulation” (ILR). Individuals live in groups of variable size, connected by random migration, with purely local competition and cooperation. After introducing this structure and quantifying group size rigididy in Section 2, we analize it in Sections 3, 4 and 5 under the assumptions that the number of groups and the typical size of the groups are both large and selection is weak. We conclude (1) When migration prevails over rigidity, the picture suggested by Hamilton is accurate, and a sharp condition for the spread of cooperation is obtained, extending Hamilton’s condition to the non-linear case. (2) When rigidity prevails over migration, we have a result identical to Taylor’s [32], with costly cooperation being selected against.

Clarifying the role of local competition on the spread of cooperation is particularly relevant because a large number of recent papers (see, e.g., [27, 8, 18, 19, 23, 24]) give a central role to the population structure of [32], as they study evolution of behavior on this structure, or introduce somewhat modified versions of that population structure and give it a benchmark status. A number of studies have neglected elasticity in modeling the evolution of cooperation and altruism under population viscosity, to avoid the technical complications that it requires (e.g., Section 7.1 of [7], and [8]). Our concern is that in so doing, these studies have suggested, even if unintentionally, that under purely local competition the Taylor regime is typical. In our analysis, the Taylor regime requires group-size rigidity to dominate migration. The task of deciding what levels of rigidity and migration are realistic should be addressed empirically, and the true range of the Hamilton and the Taylor regimes in natural populations remains to be assessed. It is possible that in many situations migration and rigidity are comparable forces, in which case we expect cooperation to spread, but under conditions that are more stringent than those given in the Hamilton regime.

Our derivation of the extension of Hamilton’s rule in Section 4 will also indicate that this “beta version of Hamilton’s condition” should be of some generality. As we will explain in Section 7, what produces this condition is a certain structure of the coalescent process associated to the population structure in the absence of selection. The relevant assortment of genes can be computed by considering a single focal group, following the lineages of its members back in time until each one migrates out of the group, and relating this coalescent and the gene frequency in the population to the gene frequency in this focal group. Since several different population structures lead to a similar coalescent process within a group, the expression of gene assortment can be seen to be fairly general. This explains, for instance, why in [29], display (3), we obtained the same beta version of Hamilton’s rule for invasion, in the context of a different population structure, in which groups compete directly among themselves. This observation allows us to apply to ILR in the Hamilton regime the consequences of display (3) in [29]. This applies, for instance, to the analysis in that paper of the conditions for invasion of altruistic contingent behavior, modeled as a trigger strategy in an iterated public goods game. We can therefore conclude also in the ILR setting, that such altruistic behavior can spread with realistically modest levels of relatedness in groups. Further analysis and applications of the beta version of Hamilton’s rule are provided in [4] and [3], showing the simplicity and relevance of some of its consequences.

## 2 Islands with local regulation

First we introduce the population structure in the absence of selection. This population structure is similar to Wright’s island model, but group size is not fixed and instead is allowed to vary around a typical value *n*_0_. The population is split into *g* groups. Generations are non-overlaping and reproduction is asexual. In the beginning of a new generation cycle, groups contain only adult individuals. The size of a group is the number *n* of these individuals. In the beginning of a cycle, adults produce offspring that remain in the group until the end of the cycle when they reach adulthood, if they survive. Each adult produces a random, statistically independent, number of adult offspring, with a distribution that depends on the number of adult individuals in its group. When group size is *n*, the mean of this individual adult offspring distribution is *h*(*s*), where *s* = *n/n*_0_ is the scaled group size, *h*(1) = 1, *h*(*s*) has a continuous derivative *h^′^*(*s*) *<* 0, and *sh*(*s*) *<* 1, when *s* < 1. Thus *h*(*s*) captures the idea that increasing group size decreases fitness. At the end of the cycle the adults present in the beginning die, while each adult born in that cycle either migrates to a randomly chosen group with probability *m*, or otherwise stays in its group of birth.

We assume that *g* is large. This implies that migration is long range and that averages across groups or individuals are well approximated by expected values for a randomly chosen focal group or individual. We also suppose, unless when stated otherwise, that *n*_0_ is large. This implies that fluctuations in size across groups can be neglected. In the absence of migration, the scaled group size behaves then as a discrete time dynamical system, with *s* being mapped to *sh*(*s*) in each iteration. The assumptions about *h*(*s*) imply that if initially *s* < 1, then *s* increases towards *s*_0_ = 1. If initially *s* > 1, then *s* decreases towards *s*_0_ = 1. The speed at which this happens depends on the derivative 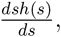 at *s* = 1. Standard computations [21] show that the relaxation time is of order 1*/λ*, where *λ* = − *h*′(1) will be called “the rigidity parameter”, and will play a major role in this paper. The assumptions on *h*(*s*) imply that 0 < *λ* ≤ 1. A small value of *λ* indicates a very elastic group environment, and a large value a very rigid environment. (The assumptions that we are making on *h*(*s*) can be somewhat relaxed, but attraction to the equilibrium at *s* = 1 could occur then through damped oscilations, rather than monotonically. We are focusing on the simpler monotonic picture for simplicity. See [21] for a detailed analysis of the dynamical system that maps *s* to *sh*(*s*), and their Table 2, for a list of forms that *h*(*s*) has taken in the literature.)

Groups are also pulled towards a common size by migration. Since migration is assumed to occur at a fixed rate, larger groups will produce more migrants than smaller ones, tending to equalize group size in a time of order 1*/m*. The comparison between the two time scales for relaxation, 1*/λ* and 1*/m*, will be the key behind our analysis.

The combined effects of rigidity and migration, will produce an equilibrium, in which each group has size close to *n*_0_. The number of migrants produced by a focal group in a cycle will then be close to a binomial with *n*_0_ attempts and probability *m* of success. The number of migrants that the same group receives in a cycle is well approximated by a Poisson distribution with mean *n*_0_*m*, because there are *gn*_0_*m* migrants, each one migrates to the focal group with probability 1*/g*, and *g* is large. The net flux of migrants into the focal group is hence a random variable with mean 0, and variance of order *O*(*n*_0_*m*). Therefore, this net flux produces a change in *s* of the order of 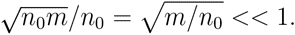

Selection is introduced by assuming that each individual is of type A or N (typically representing cooperators and non-cooperators), and that the type is inherited by the offspring without mutation. The expected number of adult offspring of an individual of type * (representing A or N), in a group with *k_A_* individuals of type A and *k_N_* individuals of type N will be supposed to be 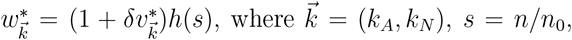 with *n* = *k_A_* + *k_N_*, the parameter *δ* ≥ 0 measures the strength of selection, and the quantities 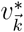 are in principle arbitrary but assumed to be bounded in absolute value by some *v*_max_. (In order for *δ* to properly account for the strength of selection, we suppose, with no loss of generality, that the order of magnitude of *v*_max_ is that of 1.) We assume that when *k_A_* = 0, we have 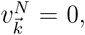 so that in case all individuals are of type N, the model behaves as the neutral one. When *δ* = 0, we also recover the neutral model, and the types A and N are then neutral markers. Selection operates on a time scale of order 1*/δ*. When *δ >* 0 is small enough that this time scale is longer than the other relevant ones, namely 1*/λ* and 1*/m*, selection is weak. In this case it is natural to refer to the quantities 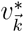 as “marginal fitness functions”. We will denote by 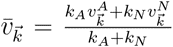 the average marginal fitness function in a group with composition 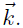 In case the interaction among group members is well represented by a linear public goods game (PG), we have 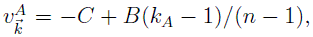 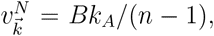 where 0 < *C* < *B* are the usual cost and benefit parameters. When *n* is large, it is natural to assume that 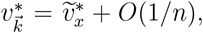 with 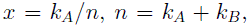 for some piecewise continuous function 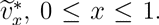 For instance, for the PG we have 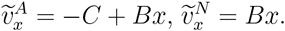

## 3 Weak selection and quasi-equilibria

In this paper we will always suppose that selection is weak, in the sense that *δ << m* and *δ << λ*. This implies that for each frequency *p* of types A, groups will reach a quasiequilibrium in a time much shorter than 1*/δ* and this lasts for a period of time of order 1*/δ*, before changes in *p* become relevant.

We can think of the quasi-equilibrium with frequency *p* of types A as a small perturbation of what we would observe if we had *δ* = 0. With *δ* = 0, we would have the neutral equilibrium described in the previous section. The fraction of type A individuals in the population would be *p*, which would vary by drift only in a time scale of order *g*^2^, assumed much larger than all the relevant time scales, including 1*/δ*. Selection with a small *δ >* 0, perturbs this picture, and leads to changes in the values of *p* and *s* in a time scale of order 1*/δ*. The direction in which *p* varies will be our main concern below. But before we can address this question, we note that the effects of weak selection on the value of *s* will be small, in the sense that at any time and for any value of *p* we will have *s* = *s_p_* = 1 + *O*(*δ/λ*). This fact is a consequence of the fact that fitnesses are always between the two values given by (1 *± δv*_max_)*h*(*s*). Therefore they are larger than 1, when *s* is below *s*_min_ that solves (1 − *δv*_max_)*h*(*s*_min_) = 1. And they are smaller than 1, when *s* is above *s*_max_ that solves (1 + *δv*_max_)*h*(*s*_max_) = 1. Solving each one of these two equations, with the notation *s*_m_ for *s*_min_, or *s*_max_, gives *h*(*s*_m_) = 1 + *O*(*δ*). Since *h*(*s*) is continuously differentiable, with *h*(1) = 1, *h^/^*(1) = *-λ*, we have *h*(*s*) = 1 *-λ*(*s-* 1)+*o*(*s-* 1) and hence *s*_m_ = 1 + *O*(*δ/λ*). To complete the argument, observe now that *s_p_* must be constrained to be inside the interval [*s*_min_, *s*_max_], since it would be pushed back up if becoming smaller than *s*_min_ and back down if becoming larger than *s*_max_.

The quasi-equilibrium with a given value of *p* can now be described as follows, thanks to a well known result by S. Wright [38] on the distribution of alleles in the infinite island model. Groups have size close to *n*_0_, and the distribution of the fraction *x* of types A over these groups is therefore well approximated by that of an infinite islands model with this group size, namely a beta distribution with parameters *lp* and *lq*, where *l* = 2*mn*_eff_ and the effective population size is *n*_eff_ = *n*_0_*/σ*^2^, where *σ*^2^ is the variance in the number of adult offspring that each individual produces in a life cycle (see [6], pp. 105,6, but be aware that his *N* relates to our *n*_0_ as *N* = 2*n*_0_). For instance, when the adult offpring distribution is Poisson, we have *n*_eff_ = *n*_0_. The parameter *l* is related to the relatedness *R* between individuals in the same group through the expression 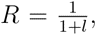 that allows one to obtain estimates of *l* from empirical estimates of *R*.

Since we will focus on a situation in which *s* = *n/n*_0_ is typically close to 1, the main way in which the function *h*(*s*) is relevant is through the value of the rigidity parameter 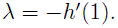 Note that the larger *λ*, the stronger is the effect of intra-group competition in decreasing fitness.

Our goal is to compute the variation Δ*p* of the value of *p* over one generation, when the population is in one of the quasi-equilibria, for this we will use the well known formula

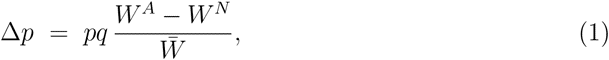

where *W ^*^* is the average number of adult offspring of individuals of type *, and 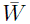 is the average number of adult offspring of all the individuals in the population.

To compute *W ^*^*, one can select a focal individual at random from the population, and compute its expected number of adult offspring, conditioned on its type being *. Now, the lineage of the focal individual should have been in its current group for a random number of generation, of order 1*/m*. That lineage produced other individuals of type * in the focal’s group. This affects the fitness of the focal individual in two ways. On one hand it produces assortment of identical types, and hence a larger fitness for a focal cooperator (viscosity promotes cooperation). On the other hand the behavior of individuals created by that lineage in the focal’s group could have had an impact on the current size of that group, and hence on the fitness of its members. A focal cooperator means a group that had more cooperators than average in the recent past, and they may have lead to an expansion of that group size which, through regulation, cancels the benefits of cooperation on a focal cooperator (viscosity not enough to promote cooperation). Whether this effect of focal’s type on group size is relevant depends on the relationship between *m* and *λ*, as we will see in the next two sections.

## 4 Hamilton regime

Suppose *λ << m*. As explained above, we suppose that the population is in a quasiequilibrium, and we select at random a focal individual, in order to compute the average fitnesses that appear in (1). The lineage of the focal individual has been in the focal’s group for a number of generations of order 1*/m*. In each of these generations the scaled groups size could only have been affected by the group composition by an amount at most of order *δ*. Over *O*(1*/m*) generations this is an amount *O*(*δ/m*). Hence the scaled size *s^*^*of the focal’s group satisfies, regardless of the type of the focal, *|s^*^ - s_p_|* = *O*(*δ/m*). And since *s_p_* is close to 1, and *h*(*s*) is continuously differentiable, with *h^/^*(1) = *-λ*, we must have *|h*(*s^*^*) − *h*(*s_p_*)*|* = *O*(*λδ/m*) = *o*(*δ*). Denoting by 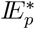 the expectation conditioned on the focal being type *\**, we obtain

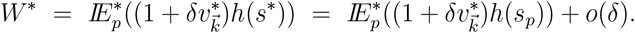

We can now apply this to (1), once with *\** = *A*, once with *\** = *N* and then note that 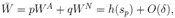, to obtain

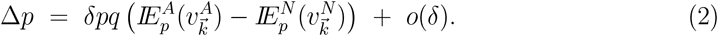

This expression can be made more explicit, by using the beta distribution described in the previous section. We will use the notation beta 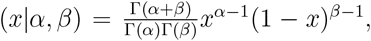 to denote the probability density of a beta distribution with parameters *α* and *β*. The only special care that needs to be taken is that conditioning on the type of the focal modifies the allele distribution in its group. For instance, the information that the focal is type A indicates that there are more types A in the group than a randomly selected group would have (sampling bias). This can be easily taken care of, using Bayes’ formula. Since given that a group has a fraction *x* of types A yields probability *x* that an individual taken at random from this group will be type A, we obtain an extra factor *x* in the density, when conditioning on the focal being type A. Normalizing the distribution, gives a beta with parameters *lp* + 1 and *lq*. Analogously, conditioning on the focal being type N, yields a beta with parameters *lp* and *lq* + 1. We can now write (2) as

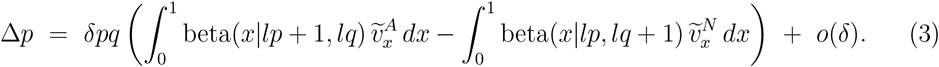

In the case of the PG, the difference between the integrals can be easily computed as − *C* + *BR*, with 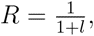 Wright’s expression for the relatedness in the infinite island model (for haploids). In this case, types A grow in frequency under Hamilton’s rule *C < BR*. In the more general case, covered by (3), this condition is replaced by the inequality among the two integrals that makes the large parentesis positive. we call this inequality “the beta version of Hamilton’s condition”. If we let *p* → 0, we obtain a condition for types A to invade when rare. This one is particularly simple, because the second integral converges then to 0, while the first one simplifies considerably, and normalizing factors can be dropped. The result is the “beta version of Hamilton’s condition for invasion”:

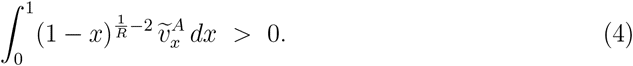

The picture that emerges in the Hamilton regime is as follows. In a quasi-equilibrium, groups are maintained at very similar sizes by migration. Types A and types N find themselves then in groups of same typical size and their adult offspring productivity depends only on the relative composition of their group, and not on its size. Assuming that cooperation is individually costly, in each group non-cooperators will be more productive than the cooperators. But groups with many cooperators will be more productive, and the strong migration will make this extra productivity translate into extra production of immigrants, rather than affect group size. Under strong enough assortment, cooperators will then in average be more productive than non-cooperators. Assortment, produced by identity by descent, will dictate the direction of selection, as quantified by (3).

## 5 Taylor regime

Now we turn to the case *m << λ*. Our goal is to complete the argument that costly cooperation is eliminated in this case. The technical definition of costly cooperation here is very mild: we suppose that 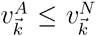 for every 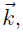 with strict inequality in at least one case. In other words, in each group, at any time, types A are not better off than types N and are sometimes worse off.

As explained in Section 3, the lineage of the focal individual must have been in the focal’s group for a time of order 1*/m >>* 1*/λ*. Now, since *δ <<* 1, the dynamics in this group during this time is a small perturbation of the dynamics with *δ* = 0, and therefore its relaxation time is close to 1*/λ*. Therefore, the evolution in this group over the time that the lineage of the focal was inside it is sufficient to have produced equilibrium in group size, meaning that it reached scaled size *s* = *s^*^* that satisfies 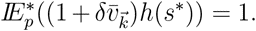 Now, the assumption of costly cooperation implies that 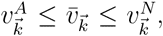 for every 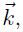 with strict inequalities in at least one case. Therefore

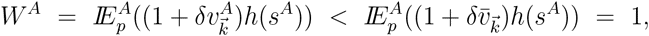

and,

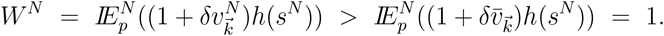

Applying these two inequalities to (1) gives

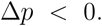

The picture that emerges in this regime is as follows. In a quasi-equilibrium, group sizes may differ significantly from each other, depending on their composition. Groups that have more cooperators will tend to be larger. Migration is not strong enough to equilibrate group size against this effect, so that cooperation produces larger groups. All groups will come to local equilibrium, with same average productivity. If cooperation is individually costly, cooperators must then have lower productivity than non-cooperators and will be eliminated, in a tragedy of the commons scenario.

## 6 A note on smaller group size

The assumption that group size is so large that its normal fluctuations can be ignored is an idealization that simplified the analysis, but it is of obvious interest to know what happens if groups are smaller. In this case, a focal group in a quasi-equilibrium, is no longer well described by a deterministic dynamical system, but rather by a Markov chain, with fluctuating group size. Still, within a range of 3 standard deviations, and with *n*_0_ even as small as 10, we should expect scaled group size to vary typically between at most 0 and 2. Under the assumption that *h^/^*(*s*) does not vary much in this range, the arguments from the previous sections can be used in good approximation.

The arguments in Section 5 continue to hold when *m << λ*. The arguments in Section 4 continue to hold when *λ << m*, but only to the point of yielding (2). The expected values in (2) do not yield then simple expressions, as was the case in (3). Still, in the case of the PG, the difference of these two expectations is again −*C* + *BR*, where *R* is the relatedness of the focal individual to a distinct randomly chosen individual from the same group. To see this, define *κ* as the fraction of types A among the members of the group in which the focal is, but excluding the focal. Then 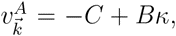 and 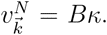 The difference in expectations in (2) is then 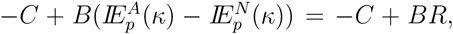 where we are using the fact that the difference in expectations here is identical to the regression coefficient of *κ* on the indicator that the focal individual is type A (see, e.g., Section 2 of the Supplementary Information of [29]). It is conceivable that for an interesting range of values of the parameters, (2) could be replaced in good approximation by (3), with the parameter *l* computed from effective population size and migration rate.

## 7 Discussion

The analysis of the ILR presented above sheds light on several aspects on how population viscosity can allow for the spread of cooperation and altruism. ILR is a population structure in which cooperation and competition are purely local. Migration is completely random and independent of local conditions. In this setting we concluded that group rigidity only poses a problem to the spread of cooperation if it is a stronger force than migration. The picture is like this: if migration is weak compared to rigidity, local cooperation leads to crowded groups in which further cooperation is of lesser importance than local competition (Taylor regime). But if migration is a stronger force than rigidity, then such overcrowding does not happen (Hamilton regime). Groups with more cooperators are then more productive and contribute more migrants to the population, and in this way cooperation spreads provided that in average cooperators have a greater marginal fitness than non-cooperators. This happens, when migration, inspite of being stronger than rigidity, is also weak enough to allow for assortment of cooperative alleles, through identity by descent. The precise expression of this assortment condition in ILR under weak selection is Hamilton’s rule in the case of additive gene action, and the more general beta version in case of possibly non-additive gene action.

We note the “non-zero-sumness” associated to the spread of cooperation in the Hamilton regime. For simplicity and concretness, consider the additive gene action case (linear public goods game, abbreviated PG) with *C < BR*. Linearity, in this example, implies that the average fitness of individuals in the whole population is given by (1+*δp*(*B − C*))*h*(*s*), where *s* is the average scaled group size. As *p* grows to 1, *s* must grow, and reach equilibrium at *s* = *s*_1_ that solves (1 + *δ*(*B − C*))*h*(*s*_1_) = 1. The solution is *s*_1_ = 1 + *δ*(*B − C*)*/λ* + *o*(*δ/λ*). If initially we had no altruists and they invaded the population, thanks, say, to rare mutations, then after they fixate, the size of the population would be larger than it was initially (by an amount of order *δ/λ*). Studies that suppose inelastic group size also assume zero-sum conditions (e.g., [27]). But in doing so they are throwing away an important component of cooperative behavior. Cooperation may increase the size of a species population, at the expense perhaps of members of other species, or at the expense of abiotic resourses.

We expect the beta version of Hamilton’s rule to be a good approximation in settings that are less idealized than ILR, or that incorporate other features, like global competition, group fissioning, and group extinction. The elements of the population structure that yielded (3) in the ILR setting have two components. First migration must be strong enough, as compared to rigidity (fitness sensitivity to group size) to imply that the fitness of cooperators and non-cooperators are in general not being affected by overcrowding. And second, that when we select a focal individual at random and analyze its lineage, we obtain a coalescent structure close to that of Wright’s islands model. In that coalescent structure, the lineage of the focal individual stays in the same group for a random number of generations that has a geometric distribution with probability *m* of success. The same happens with the lineage of other individuals in the same group, and coalescence happens when lineages come together. From generation to generation, moving into the past, individuals take their progenitor at random in the group, independently, so that coalescence between two lineages happens with probability given by inverse of group size. This coalescent structure yields the beta distribution of alleles (see, e.g., [36]). Clearly, in ILR with large *n*_0_ we have a coalescent structure that is close to this description. But now modify ILR by adding group extinction with probability *φ* in each generation. If *φ << m* and *φ << λ*, then the coalescent structure is not severely affected (regardless of the way recolonization is assumed to happen). Therefore, if also *δ << λ << m*, then the conditions for the spread of cooperation would be the same ones obtained in Section 4. If, on the other hand, *φ* is comparable to or larger than *m* or *λ*, then a modified coalescent has to be analyzed to decide the fate of the cooperative alleles.

These observations explain what would otherwise have seemed as a surprising coincidence. In display (3) of [29], we obtained the same condition for invasion (4) for a population structure that is substantially different from ILR. That population structure was called there “two-level Fisher-Wright with selection and migration” (2lFW). The idea in that population structure goes back at least to [9], who proposed it as a way for viscosity to promote the evolution of cooperation, without the hindrance of local competition. This population structure was analyzed under weak selection for the PG in [20] and [8], and called “budding viscosity model” in the latter paper. In 2lFW we again have a population of haploid individuals, divided into many groups, with asexual reproduction and non-overlaping generations. Groups have a fixed size *n*, and compete directly with each other, in that in each new generation, each group has a progenitor group chosen from among all groups in the previous generation independently, with probabilities proportional to average group fitness among the candidate groups. Individuals in each group in the new generation then chose their parents from among the members of the progenitor group, independently, with probabilities proportional to individual fitness. This form of group competition allows for average group fitness to be different from 1, even if groups have constant size. (In contrast, in ILR the group size flexibility is needed for this to be possible. Note that 2lFW is a structure with global competition, in contrast with ILR). In the neutral version of 2lFW, groups choose their progenitor group uniformly over the previous generation, and then individuals in each group choose their parent uniformly over the members of the progenitor group. When analyzing gene assortment in the neutral version of 2lFW, using the coalescent, one selects a focal individual and follows its lineage back in time. The group to which this individual belongs will be changing as we move back in time, but this has no consequence. Since all the individuals in a group in each generation choose their parent from the same progenitor group, we have exactly the same coalescent as in Wright’s islands model. This explains the coincidence in the condition (4) for ILR and 2lFW, and also suggests again that while not completely universal, this condition for invasion of a cooperative allele has a very broad range of validity, at least as a good approximation. (This reasoning shows that also (3) will hold in the 2lFW setting, when *δ << m*, *n* is large and 2 *nm* = *l*.)

Real populations are more complex then ILR and 2lFW in many ways. Competition typically will be partly within the group and partly with other groups, if, for instance, resources are in part obtained from a territory common to many groups, or when groups move in search of better foraging grounds. In these cases, the regulation function is better described as *h*(*s, s_e_*), where *s_e_* = *n_e_/n_e_*_0_ is the scaled population size of the “economic neighborhood” (possibly the whole population) of a focal group, assumed to have typical population size *n_e_*_0_ and current population size *n_e_*. Our analysis extends to this more general setting, with the rigidity parameter now defined as 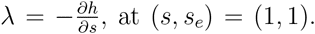 Competition that is partially non-local should yield values of *λ* smaller than in the purely local case, discussed before. Groups may also go extinct, or split or produce propagules that generate new groups. In such cases, group size may often be different from the typical *n*_0_. But if extinction and spliting occur infrequently and fitness is substantially higher than 1 in groups that are significantly smaller than *n*_0_, then this would not be the case, as small groups then grow to sizes comparable to *n*_0_ in a few generations. The arguments in the previous two paragraphs, indicate that under the scenarios discussed here, the beta version of Hamilton rule should be a good approximation when *δ << λ << m*, since the coalescent is then always close to that of the infinite islands model.

This discussion suggests that the beta version of Hamilton’s rule may be a good approximation under very broad conditions. But if so we are faced with the task of estimating *λ* for real populations to compare it to the migration rate. Unfortunately, this is a very difficult and controversial task, as explained for instance in [35] and [30].

One pivotal observation is the tension that exists between two conditions needed for the spread of costly cooperation and altruism through population viscosity, under local competition. On one hand *m* must be large compared to *λ*, to avoid the overcrowding of the Taylor regime. On the other hand, a large *m* reduces assortment (in particular reduces relatedness), making the conditions for invasion of cooperation (say, *C < BR* in the case of PG) harder to fulfill. Of course, a very large *B/C* ratio in PG would provide a solution. There are empirical cases in which indeed the benefits seem to be much larger than the costs [26], and it is important to investigate empirically what values of *B/C*, *m*, *n*_0_ and *λ* are reasonable to expect, and whether they are compatible with the spread of altruism under additive gene action across group members (PG). Here we want to repeat now a central message from [29] and point out that even if data may show that typically *C > BR*, contingency of cooperative/altruistic behavior may be a key to its spread. This was done in that paper in the context of 2lFW, and we can now extend the conclusion to ILR and local regulation. In that paper we considered a version of the iterated linear public goods game (IPG) introduced in [16] and [2]. Suppose that a linear public goods game is played *T* times in a life cycle, cooperators cooperate in the first round and later only if their payoff in the first round was positive. The scaled marginal fitness function for cooperators takes then the form 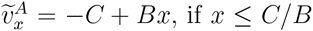 and 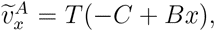 if *x > C/B*. The analysis in [29] shows that when *T* is large and *C/B* is small (4) yields in good approximation the simple condition *C < BR* ln *T* . This provides a substantially larger range of values for *B/C*, *m*, *n*_0_ and *λ* than if cooperators acted unconditionally, in which case the condition for invasion is simply *CT < BT R*, which simplifies to the usual *C < BR*. For instance, if *B/C* = 4, *m* = 0.1, *n*_0_ = 100, *λ* = 0.01, then *λ << m* is satisfied, and, assuming *n*_eff_ = *n*_0_, as in the case of Poisson adult offspring distribution, we have *R* = 1*/*(1 + 2*n*_0_*m*) = 0.048 (comparable to commonly found values, [15, 11]). The condition for invasion becomes here *T > e^C/^*^(^*^BR^*^)^ = 190. As we argued in [29], for some species including humans, certain cooperative activities, including hunting in large groups, and food sharing are potentially repeated thousands of times in a life-cycle, so that a value of that order of magnitude is quite realistic. (If cooperators only give up cooperating after a certain number *a* of attempts that resulted in negative payoff to them, then the parameter *T* will be the number of games potentially played in a life-cycle, divided by *a*. If we assume *a* = 5, then *T* = 190 corresponds to 950 potential repetitions of the game.)

Needless to say, in many real populations some of the assumptions required above to obtain (3) will not hold. If groups split often and grow slowly after splitting, if extinction with recolonization happens frequently, if migration is short ranged, if groups are small, and in several other cases, the conditions for the proliferation of a cooperative allele may not be well approximated by the beta version of Hamilton’s rule. But even then, the methods from this paper suggest a line of investigation based on the analysis of coalescent structures and on how much they are affected by the type of the focal allele. As we observed in Section 6, in some situations (2) may still hold, even if the expectations in these expressions may not be well approximated by expectations under beta distributions. The remarks in this discussion section suggest also that the beta version of Hamilton’s rule should be a good point of reference from which one can start a deeper investigation on how gene assortment in viscous group structured populations allows the spread of cooperative and altruistic alleles.

## Acknowledgments

We are grateful to Clark Barrett, Maciek Chudek, Sarah Mathew and Karthik Panchanathan for enlightening discussions. Partially supported by CNPq, under grant 480476/2009-8.

